# Cancer-associated recurrent mutations in RNase III domains of DICER1

**DOI:** 10.1101/005686

**Authors:** Bülent Arman Aksoy, Anders Jacobsen, Robert J. Fieldhouse, William Lee, Emek Demir, Giovanni Ciriello, Nikolaus Schultz, Debora S. Marks, Chris Sander

## Abstract

Mutations in the RNase IIIb domain of DICER1 are known to disrupt processing of 5p-strand pre-miRNAs and these mutations have previously been associated with cancer. Using data from the Cancer Genome Atlas project, we show that these mutations are recurrent across four cancer types and that a previously uncharacterized recurrent mutation in the adjacent RNase IIIa domain also disrupts 5p-strand miRNA processing. Analysis of the downstream effects of the resulting imbalance 5p/3p shows a statistically significant effect on the expression of mRNAs targeted by major conserved miRNA families. In summary, these mutations in DICER1 lead to an imbalance in miRNA strands, which has an effect on mRNA transcript levels that appear to contribute to the oncogenesis.

## Brief Communication

MicroRNAs (miRNAs) are small non-coding RNA molecules that regulate expression of their transcript targets [1] DICER1 is a key enzyme that is responsible for cutting the 5p and 3p strands of the pre-miRNA in the early stages of the miRNA biogenesis. Processing of the 5p and 3p strands, which is carried out by the RNase III domains of DICER1, is necessary for loading the functional miRNA strand into the RISC complex. Previous studies have identified recurrent mutations in the RNase IIIb domain in different cancer types [2, 3, 4, 5, 6, 7, 8, 9]. These mutations (at residiues E1813, D1810, D1709, E1705 and R1703) were shown to be in the active site of the enzyme and were proven to disrupt the processing of the 5p stand of the miRNA [10]. Others have shown that hotspot mutations in the RNase IIIb domain cause depletion of 5p strands relative to their corresponding 3p strands, leading to an asymmetry in the abundance of the two [11, 7].

Although the asymmetry in the miRNA processing due to hotspot mutations has been characterized using model organisms; the effect of this miRNA depletion on the mRNA levels have not been studied extensively in the context of the human tumors. It is, for example, unknown whether it is the 5p-strand depletion or increased 3p-strand accessibility that promotes the cancer. In either of the cases, it is also unknown whether there is any particular miRNA or miRNA family of which depletion or over-expression drives this phenotype. In this study, using human tumor data from the Cancer Genome Atlas (TCGA) project, we wanted to better characterize the effects of *DICER1* mutations on miRNA and mRNA profiles of the patients.

We first asked whether we could observe the asymmetry in the miRNA processing using the miRNA-Seq data. For this, we looked whether any of the previously identified hotspot mutations were present in the TCGA data set (14 cancer types, 5535 sequenced samples). We found that 15 out of 123 *DICER1* mutants carried a mutation in the RNase IIIb domain of the protein at a previously identified hotspot (Figure 1a). After filtering out cases that were hyper-mutated and samples that did not have miRNA-Seq data available, we were left with 8 *DICER1* hotspot mutants. We then compared the miRNA levels in these hotspot mutants to the miRNA levels in 3171 *DICER1* wildtype tumors across multiple cancers. Confirming the results of the previous studies, we saw 5p strand miRNAs were relatively down-regulated in mutants and the changes in the expression of 5p strands were significantly different than the 3p strands (Wilcoxon rank sum test; *p* < 10^−29^; Figure 1b-c).

**Figure 1:**
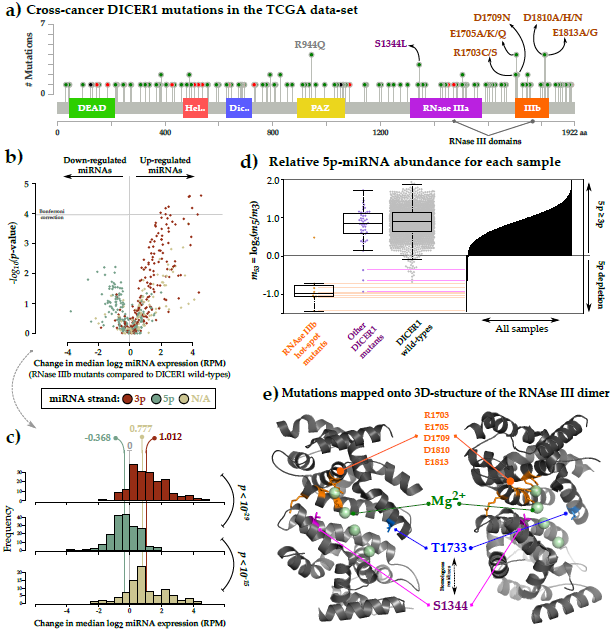
Disabling mutations in RNase III domains of DICER1 lead to 5p miRNA depletion in cancer. **a)** A majority of the hotspot mutations in the RNase III domains of the *DICER1* are present in the Cancer Genome Atlas project across multiple cancer types. **b-c)** Hotspot mutations in the RNase IIIb domain cause relative down-regulation of 5p-stand and up-regulation of 3p strand miRNAs in mutants compared to DICER1 wild-types. **d)** Hotspot mutated samples tend to have relatively lower 5p miRNA abundance compared to *DICER1* wild-type cases. Using sample-specific relative 5p abundances, we identified three more DICER1 mutated cases that also show 5p-depletion phenotype (*m*_5,3_ *<* 0). **e)** Two out of three cases, who has relatively low 5p abundance, had a S1344 mutation in the RNase IIIa domain that is responsible for processing the 3p strand of the miRNA. The mutated amino acid, S1344 in RNase IIIa domain, is homologous to T1733 in RNase IIIb domain, which in turn is evolutionary coupled to the hotspot mutations. This indicates that S1344, although it is in RNase IIIa domain, is important for proper functioning of the RNase IIIb domain.

Having observed a phenotype characterized by relative 5p strand depletion in hotspot RNase IIIb mutants, we asked whether any of the other *DICER1* mutants had a similar phenotype. To investigate this, we first estimated the abundance of 5p strands relative to 3p strands for each patient: 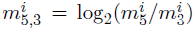, where 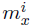 is the median expression of the *x*-strand miRNAs in patient *i*. As expected, the majority of the hotspot mutants had exceptionally low 5p-strand abundance compared to *DICER1* wildtypes (Figure 1d).

In addition to the known hotspots mutants, we identified three more *DICER1* mutant cases that had relatively low 5p abundance 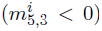. One of these three *DICER1* mutants had a hotspot mutation in its RNase IIIb domain, but was excluded from the initial analysis because it was a in hyper-mutated sample (Table S2). Surprisingly, the other two cases with low 5p abundance had an S1344L mutation in the RNase IIIa domain that is responsible for processing the 3p strand of the miRNA.

As the observation of recurrent mutations in cancer samples is consistent with a selective functional impact of the mutation, the question arises as to the effect of the S1344L mutations on the catalytic function of the RNase domains. Inspection of the 3D structure (or model) of the individual domain reveals that residue S1344L (in domain IIIa) and its homologous residue T1733 (in domain IIIb) are far from the active site residues (19.60±2.62Å distance) in their respective domains (Figure 1e). However, evolutionary couplings [12] between S1344L/T1733 and the active site residues, as deduced from co-evolution patterns in the multiple sequence alignment of RNase III-like domains, are fairly strong. The contradiction is resolved by inspection of the model of the RNase IIIa - IIIb heterodimer (as inferred from the crystal structure of the RNase IIIb homodimer) [10]. In the heterodimer, S1344L in domain IIIa is close (11.72±1.98Å distance) to active site of domain IIIb (residues E1813, D1810, D1709, E1705 and R1703) and T1733 in domain IIIb is close to the active site residues of domain IIIa. These residue arrangements and functional couplings are beautifully consistent with the observation that mutations in S1344L in domain IIIa affect 5p processing, as observed in our analysis of the effect of these mutations on the balance of 3p/5p miRNA expression profiles in cancer samples. This is consistent with the earlier observations that mutations in the active site residues of domain IIIa affect 3p processing, while mutations in the active site residues of domain IIIb affect 5p processing. The subtly of the difference between the earlier and current observation lies in the residue interactions across the heterodimer interface [13] and in fact the earlier observation of 3p/5p asymmetry are confirmed here by completely independent observation in human cancer samples.

Other studies have shown that *DICER1* hotspot mutations are biallelic in cancer, where a disabling mutation acts as the second hit to the enzyme [5, 6, 8] Based on this observation, the relative 5p depletion phenotype of RNase III mutants in our analysis suggested that these patients also had a second event disabling the other *DICER1* allele. To address this question, we re-analyzed the sequencing data available for *DICER1* mutant cases, this time using a different pipeline that can better identify insertions or deletions. In a majority of the *DICER1* RNase III hotspot mutant samples, we were able to identify a secondary disabling genomic event affecting the other *DICER1* allele (Table S3). Furthermore, we found that these biallelic mutated cases had lower 5p abundance than the other *DICER1* mutants in our earlier analysis.

Having identified possibly functional mutations in *DICER1* and their effect on the miRNA profiles, we tested whether these mutations lead to functional changes in the mRNA profiles. Others have previously characterized *DICER1* hotspot mutations using mouse-derived cell lines as *in vitro* models [7, 8, 11] These studies have shown that the mRNA profiles of cell lines with different DICER1 RNase IIIb hotspot mutations had different mRNA signatures compared to the *DICER1*-wildtype cell lines. They further found an association between the down-regulated miRNAs and their differentially-expressed target transcripts, which suggests a differential regulation of the mRNA levels due to asymmetric miRNA processing in *DICER1* hotspot mutants.

Although there is *in vitro* evidence that the asymmetry in the miRNA processing lead to significant changes in the mRNA profiles; there are no previous reports that describe the differential mRNA expression in accordance with the miRNA expression data from human tumors. To this end, we identified 12 cases across four cancer types that both had RNA-Seq data available and carried a hotspot RNase III mutation either in the IIIa or IIIb domains of the DICER1 protein. We then wanted to check whether we could identify a common mRNA expression signature for these *DICER1* RNase III hotspot mutants in comparison to 1212 *DICER1* wildtype cases in those four cancer studies. For this, we decided to restrict our analysis to the Uterine Corpus Endometrial Carcinoma (UCEC) study where the RNA-Seq data set contained 8 *DICER1* RNase III mutants and 222 *DICER1* wild-types. We found 10 genes to be significantly up-regulated and none to be down-regulated in the hotspot mutated cases when compared to wildtypes (*p* < 0.05 after Bonferroni correction; Table S4). Notably, we found higher expression of *HMGA2*, a well-known oncogene and target of *let-7* miRNA family, in mutants [14, 15, 1].

Following up on this, we asked whether the up-regulated genes in mutants were targets of particular miRNA families. To answer this question, we conducted a gene set enrichment analysis (GSEA) using well-known biological pathways and well-conserved miRNA family target genes as our query gene sets [16]. Our analysis showed strong enrichment of both *let-7/98/4458/4500* and *miR-17/17-5p/20ab/20b-5p/93/106ab/427/518a-3p/519d* target genes in RNase III mutants (Table 1; FDR < %10). For both families, 5p strand of the miRNA is the predominant strand and as expected, in RNase III mutant cases, 5p-strand miRNAs that belong to these families were relatively down-regulated. Results from the GSEA also suggested that there was relatively weaker enrichment for other miRNA families and NOTCH-related pathways (Table 1; FDR < %15). A majority of the enriched gene sets (5 out of 7) represented miRNA family targets, which suggests the gene expression signature associated with these RNase III hotspot mutants is more likely to be mediated by depleted miRNA families rather than a common biological pathway. In accordance with the 5p strand depletion phenotype, a majority of these miRNA families (3 out of 5) were 5p-strand dominated. For the other two families, *miR-29abcd* and *miR-101/101ab*, although 3p is the pre-dominant miRNA strand, we saw that members of these families were down-regulated as a family in *DICER1* mutants compared to wildtype, which might be due to an indirect regulatory effect of 5p miRNA depletion.

**Table 1:** Gene sets representing targets of conserved miRNA families are up-regulated in DICER1 RNase III mutants compared to wild-types. To see the effect of relative depletion of 5p miRNAs on the mRNA profiles, we conducted a Gene Set Enrichment Analysis (GSEA) on mRNA profiles of Uterine Corpus Endometrial Cancer (UCEC) samples. We showed that targets of the major miRNA families, which are predominantly 5p-originating, are differentially up-regulated in *DICER1* mutants compared to wild-types. For each of these miRNA families, we saw consistent down-regulation of 5p strand (green) and up-regulation of 3p strand (red) miRNA members. *mut*: *DICER1* hotspot mutant; *wt*: *DICER1* wildtype; *Diff. Exp.*: Differential expression (*log2* ratio of mRNA/miRNA levels); *p value*: The probability for the null hypothesis that the genes in the set are not differentially up-regulated in mutants compared to wildtypes; *FDR*: *p* value corrected for multiple hypothesis testing.

| miRNA set | miRNA target set<br>(# of genes) | *p* value | FDR | Diff. Exp. of mRNAs<br>(8 mut.s vs 222 wt.s) | Diff. Exp. of miRNAs<br>(5 mut.s vs 107 wt.s) |
| --- | --- | --- | --- | --- | --- |
| let-7/98/4458/4500 | 90 | 0.0001 | 0.08 |  |  |
| miR-17/17-5p/20ab/20b-5p/93/-106ab/427/518a-3p/519d | 42 | 0.0002 | 0.08 |  |  |
| miR-29abcd | 87 | 0.001 | 0.11 |  |  |
| Pre-NOTCH transcription and translation | 27 | 0.001 | 0.11 |  |  |
| miR-101/101ab | 26 | 0.001 | 0.11 |  |  |
| miR-15abc/16/16abc/195/-322/424/497/1907 | 51 | 0.001 | 0.11 |  |  |
| NOTCH HLH transcription pathway | 12 | 0.002 | 0.13 |  |  |
| All | 16358 |  |  |  |  |

In summary, we showed that biallelic *DICER1* RNase III hotspot mutations, although infrequent across cancers, lead to relative depletion of 5p stand of miRNAs. In addition to known hotspot mutations, we were able to identify a previously unknown recurrent DICER1 mutation, S1344, that also leads to the 5p depletion phenotype. In accordance with the miRNA depletion phenotype, we saw up-regulation of genes that are well-known targets of the 5p-dominant miRNA families in mutant samples. It still remains unclear whether up-regulation of a particular gene, such as *HMGA2*, or activation of a particular pathway, such as NOTCH, is contributing to the oncogenesis as a result of the 5p miRNA depletion in these cells.

## Acknowledgments

We would like to thank Kjong Lehmann, Andre Kahles, Gunnar Rätsch, Özgün Babur, Pınar Aksoy, Ed Reznik, Nils Weinhold, Ruomu Jiang, Berkin Elvan for helpful discussions on the manuscript. This work was supported by US National Cancer Institute funding of the TCGA Genome Data Analysis Center (U24 CA143840).

## S1 Online Methods

The code for analyses conducted in this study and supplemental results for each of the analyses are available at http://bit.ly/dicer5p. In this study, we used miRNA, RNA-Seq and sequencing data from 14 TCGA cancer studies (Table S1).

**Table S1:** We analyzed a total of 2855 samples with miRNA and sequencing data across 14 cancer studies from the Cancer Genome Atlas.

| Abbreviation | Cancer study name | # of samples |
| --- | --- | --- |
| BLCA | Bladder urothelial carcinoma | 137 |
| BRCA | Breast invasive carcinoma | 190 |
| COADREAD | Colorectal adenocarcinoma | 241 |
| GBM | Glioblastoma multiforme | 248 |
| HNSC | Head and Neck squamous cell carcinoma | 267 |
| KICH | Kidney chromophobe | 64 |
| KIRC | Kidney renal clear cell carcinoma | 184 |
| LGG | Brain lower grade glioma | 286 |
| LUAD | Lung adenocarcinoma | 180 |
| LUSC | Lung squamous cell carcinoma | 51 |
| PRAD | Prostate adenocarcinoma | 248 |
| STAD | Stomach adenocarcinoma | 244 |
| THCA | Thyroid carcinoma | 399 |
| UCEC | Uterine corpus endometrial carcinoma | 116 |
| Total |  | 2855 |

### S1.1 Identification of *DICER1* hotspot mutations

We first asked whether previously identified *DICER1* hotspot mutations at residues E1813, D1810, D1709, E1705 and R1703 are present in TCGA data sets. For this, we conducted a cross-cancer query on cBioPortal [17] and found 123 out of 5535 sequenced samples to be *DICER1* mutated (Figure 1a and File *all_tcga-dicer1-2014_03_20.maf*). Of these 123, 12 tumor samples had at least one hotspot *DICER1* mutation in the RNase IIIb domain.

### S1.2 Analysis of the miRNA-Seq data

We next wanted to see if hotpost mutant tumors had a distinct miRNA expression profile compared to other samples. To address this question, we first obtained normalized miRNA-Seq data sets (Level 4) from the most recent TCGA analysis runs (January 15, 2014) as generated with the Firehose analysis pipeline. miRNA-Seq data for Glioblastoma Multiforme cancer study was not available from this resource, therefore, for GBM, TCGA Level 1 microarray expression data were processed and normalized using the *AgiMicroRna* R package and using settings further explained in a previous study [18, 19].

We then wanted to see whether particular miRNAs were differentially expressed in *DICER1* RNase IIIb mutants compared to *DICER1* wild-type cases. We initially excluded hotspot mutants from the analysis if they were either categorized as hyper-or ultra-mutated, or if the predicted effect of the mutation was not high as assigned by the Mutation Assesor (Table S2) [20]. To check for differential expression, we compared distribution of each miRNA expression in mutants versus wildtypes by using a Wilcoxon rank sum test. We adjusted the *p*-values using a Bonferroni correction for multiple hypothesis testing. To estimate the change in expression, we calculated the difference in median *log2*-based expression values between mutant and wildtype samples (Figure 1b-c).

**Table S2:** To identify the miRNA expression signature associated with hotspot *DICER1* mutations, we excluded hyper-mutated cases from the initial analysis. Ultra-or hyper-mutated cases tend to have higher number of somatic mutations compared to other samples. To identify miRNA profiles associated with the hotspot DICER1 mutants in a restrict way, we first conducted the differential miRNA expression analysis only on samples with relatively low number of somatic mutations (*n* < 1000).

| Sample identification | Reason for exclusion |
| --- | --- |
| TCGA-A6-6141 | Hyper-mutated sample |
| TCGA-AP-A0LM | Low allele frequency and ultra-mutated sample |
| TCGA-BS-A0UV | Low FIS and ultra-mutated sample |
| TCGA-CG-5733 | Low FIS and hyper-mutated sample |
| TCGA-D1-A17Q | Ultra-mutated sample |

To check whether the distribution of differential miRNA expression was different for different strands of the miRNA, we conducted pairwise comparisons of the differential expression values for different strands of miRNA: 5p, 3p and N/A where N/A means no strand information was available for that miRNA. For this comparison, we utilized Wilcoxon rank sum test and adjusted the *p*-values using a Bonferroni correction.

### S1.3 Additional mutation calling for *DICER1* hotspot mutants

Having observed different levels of respective 5p strand depletion in hotspot *DICER1* mutants, we wanted to see if patients with extreme phenotypes had any additional germline or somatic mutations affecting the other DICER1 allele. We, therefore, downloaded whole-exome binary sequence alignment and mapping (BAM) files for normal and tumor samples corresponding to the hotspot DICER1 mutated cases from CGHub. We then used *HaplotypeCaller* utility from the Genome Analysis Toolkit to do the joint variant calling on these BAM files [21]. To annotate the variants, we used Mutation Assesor and Oncotator tools [20].

We next used the annotated mutation file to look for new mutations that were not called by the TCGA pipeline (File: *muts_tcga-dicer1-secondcall-2014_04_09.maf*). In addition to the previously called hotspot mutations, we were able to identify other disabling *DICER1* alterations in samples that showed relatively low 5p strand abundance (Table S3).

### S1.4 Identification of evolutionary couplings in RNase III domain

In our miRNA expression analysis, in which we estimated the relative 5p strand abundance for each patient, we saw that two samples that have the biallelic S1344L mutation had considerably low 5p abundance. Based on the fact that RNase III dimerization is necessary for proper DICER1 functioning, we wanted to see how S1344L could affect 5p miRNA processing [13]. For this we ran evolutionary couplings (ECs) analysis with default settings on the EVFold server (v1.11) [12]. We provided DICER_HUMAN (UniProt:Q9UPY3) as the input protein, residues 1423-1922 of DICER1 as the sequence of interest to center the RNase IIIb domain and PDB:2eb1 as the reference structure [10]. We set the *e*-value for jackhmmer as 10^−10^ and the inference method for determining the evolutionary couplings as Pseudo Likehood Maximization (PLM).

The analysis showed that the most strongly constrained residues (with strong couplings to other residues) were 1708, 1709, 1813, 1705 and 1704. The contact maps were fairly structured, indicating they were of reasonable quality (File: *EvCouplings_DICER1_RNaseIIIb_with_2eb1.zip*). Well-known active site residues with relatively high EC strength included 1709, 1813 and 1705. We found that residues 1709, 1813 and 1705 were coupled to 1733. These ECs, however, were not consistent with the known structural constraints as 1709, 1813 and 1705 were not in close proximity to 1733 in the 3D structure (19.60±2.62Å distance).

**Table S3:** Hotspot *DICER1* mutations that lead to 5p depletion phenotype are biallelic in TCGA samples. For the majority of the hotspot *DICER1* mutants, we were able to identify a second genomic event that affect the other *DICER1* allele. These biallelic mutated samples were enriched for stronger 5p depletion phenotype (i.e. lower *m*_5,3_) compared to monoallelic alterations. *THCA*: Thyroid carcinoma; *UCEC*: Uterine corpus endometrial carcinoma; *GBM*: Glioblastoma multiforme; *COADREAD*: Colorectal adenocarcinoma; *CNA*: Copy number alteration; *HetLoss*: Heterozygous loss; *N/A*: Not available.

| Sample identifier | Cancer study | Mutation | CNA | $m_{5,3}$ |
| --- | --- | --- | --- | --- |
| TCGA-EL-A3GO | THCA | D1810H, K376fs | - | -1.43 |
| TCGA-D1-A15Z | UCEC | D1810A, L539fs | - | -1.08 |
| TCGA-EL-A3D5 | THCA | E1813G, L81fs | - | -1.05 |
| TCGA-DI-A0WH | UCEC | D1709N, M1821I, K1486fs | - | -1.02 |
| TCGA-06-2569 | GBM | E1705Q, CLPSIL1053del | Gain | -0.93 |
| TCGA-A5-A0GN | UCEC | S1344L | HetLoss | -0.92 |
| TCGA-14-0871 | GBM | Homozygous E1813G | - | -0.83 |
| TCGA-A6-6652 | COADREAD | D1810N | HetLoss | -0.71 |
| TCGA-B5-A11U | UCEC | S1344L, P1377fs | - | -0.63 |
| TCGA-D1-A17Q | UCEC | E1705K, H341P | - | -0.36 |
| TCGA-AP-A0LM | UCEC | E1705A, R490H, F1650C | - | 0.36 |
| TCGA-DM-A28C | COADREAD | E1705Q | - | 0.48 |
| TCGA-A5-A0GH | UCEC | E1813G, V1731fs | - | N/A |
| TCGA-BG-A0M6 | UCEC | E1813A | - | N/A |
| TCGA-D1-A0ZP | UCEC | R1703C | - | N/A |

A multi-alignment involving both RNase IIIa and IIIb domains indicated that S1344 in RNase IIIa domain was homologous to 1733 in RNase IIIb domain. We then inspected the corresponding locations of these residues in the 3D protein structure and found that ECs from residues 1709, 1813 and 1705 to 1733 were better explained in the RNase IIIb dimer context, where active site residues in one domain were closer (11.72±1.98Å distance) to the 1733 (i.e. S1344) in the other domain. Based on these observations, we concluded that these couplings might indicate an important role for S1344, together with other active site residues (1709, 1813, 1705) in RNase IIIb domain, in 5p strand processing.

### S1.5 Analysis of the RNA-Seq data

We next asked whether *DICER1* hotspot mutants had distinct gene expression profiles compared to other samples. To answer this question, similar to miRNA data, we obtained processed and normalized RNA-Seq data sets (Level 4) from the most recent TCGA analysis runs (January 15, 2014) as generated with the Firehose analysis pipeline. We found that THCA, GBM, COADREAD studies had RNA-Seq data for less three hotspot mutants, hindering a statistically robust comparison. We, therefore, decided to restrict our analysis to only UCEC study, where there were 8 *DICER1* hotspot mutant and 222 *DICER1* wildtype samples.

We then conducted a differential gene expression analysis using the *limma voom* R package on the gene-level RSEM counts for UCEC study and contrasted the hotspot mutant to wildtype samples [22]. We found 9 genes to be significantly up-regulated–and none down-regulated–in mutants (*p* < 0.05 after Bonferroni correction; Table S4; File: *DGE-UCEC-muts_vs_wts-allGenes.tsv*).

**Table S4:** A differential gene expression analysis comparing *DICER1* hotspot mutants to wildtypes showed 9 significantly up-regulated genes in mutants. We compared the gene expression levels in 8 DICER1 mutants to the levels in 222 *DICER1* wildtypes using the *limma voom* toolkit. We used Bonferroni correction to adjust our *p*-values for multiple hypothesis testing and found 9 genes to be differentially up-regulated in mutants (*p*_*adj*_ < 0.05). *logFC:* change in gene expression (log based)

| Gene | Gene ID | logFC | $p$-value | adjusted $p$-value |
| --- | --- | --- | --- | --- |
| HMGA2 | 8091 | 3.708 | 0.0000000001 | 0.0000016619 |
| IGDCC3 | 9543 | 3.648 | 0.0000000025 | 0.0000409144 |
| ACVR2B | 93 | 1.211 | 0.0000000083 | 0.0001365400 |
| MMP16 | 4325 | 2.333 | 0.0000002521 | 0.0041232946 |
| C17orf63 | 55731 | 0.782 | 0.0000002798 | 0.0045772958 |
| ADAMTS7 | 11173 | 1.993 | 0.0000007622 | 0.0124675442 |
| IGF2BP2 | 10644 | 3.294 | 0.0000015289 | 0.0250102395 |
| FAM171B | 165215 | 1.801 | 0.0000021387 | 0.0349852741 |
| MGAT5B | 146664 | 2.875 | 0.0000023541 | 0.0385090592 |

### S1.6 Gene set enrichment analysis (GSEA)

Having observed up-regulated genes in *DICER1* hotspot cases compared to wildtypes, we wanted to see whether these genes were targets of particular miRNAs or members of canonical pathways. To answer this question, we utilized a gene set enrichment analysis (GSEA) using the UCEC data set.

To create gene sets for targets of the well-conserved miRNA families, we first downloaded predicted miRNA targets from TargetScan (Release 6.2) and then aggregated these predictions using miRNA family-member associations to obtain a list of targets for each miRNA family [23]. We next filtered out predictions with conservation score lower than 90% and then collected targets that were in the upper 5 percentile considering their context score (i.e. scores lower than 0.3555). Using these filtered predictions, we created gene sets that were compatible with the conventional GSEA analysis [16].

We combined these miRNA target gene sets with gene sets representing well-known and curated Reactome pathways from MSigDB [24, 25]. This gave us a total of 719 gene sets, consisting of 674 gene sets for pathways and 45 for targets of miRNA families (File: *GSEA-GeneSymbols-mirFamilies_and_Pathways.gmt*). For the GSEA, we utilized the *romer* utility from the *limma* toolkit and used the contrast model that we used in the RNA-Seq data analysis [26]. We set the number of rotations to 10,000 and for each gene set, tested whether the genes in the set were enriched for any direction (up-or down-regulation).

We found genes in 7 different sets to be significantly enriched towards up-regulation and none in the reverse direction (*FDR <* 0.15; Table 1; File: *GSEA-UCEC-muts_vs_wts.tsv*). 5 out of 7 gene sets were representing target genes for miRNA families and 3 of these were miRNA families for which 5p strand was the predominant strand according to miRBase [27].

